# A simplified workflow for the analysis of whole-genome sequencing data from *Pristionchus pacificus* mutant lines

**DOI:** 10.1101/2020.11.12.379388

**Authors:** Christian Rödelsperger

## Abstract

Nematodes are attractive model systems to understand the genetic basis of various biological processes ranging from development to complex behaviors. In particular, mutagenesis experiments combined with whole-genome sequencing has been proven as one of the most effective methods to identify core players of multiple biological pathways. To enable experimentalists to apply such integrative genetic and bioinformatic analysis in the case of the satellite model organism *Pristionchus pacificus*, I present a simplified workflow for the analysis of whole-genome data from mutant lines and corresponding mapping panels. Individual components are based on well-maintained and widely used software packages and are extended by 50 lines of code for the analysis and visualization of allele frequencies. The effectiveness of this workflow is demonstrated by an application to recently generated data of a *P. pacificus* mutant line, where it reduced the number of candidate mutations from an initial set of 3,500 single nucleotide variants to ten.

## Introduction

Free-living nematodes such as *Caenorhabditis elegans* and *Pristionchus pacificus* are powerful model systems to dissect the genetic architecture of various traits and to perform comparative studies across the whole biological spectrum ranging from development to neurobiology and behavior (Sommer 2009; Sternberg 2015; Moreno *et al*. 2017; Hong *et al*. 2019). The potential of nematode systems for comparative biology was even further enhanced with the advent of high-throughput sequencing and genome editing (Lo *et al*. 2013; Nakayama *et al*. 2020). However, while reverse genetic approaches based on candidate genes are promising tools to investigate the evolution of gene function (Markov *et al*. 2016; Sieriebriennikov *et al*. 2017; Okumura *et al*. 2017), forward genetic studies in non-classical model organisms such as *P. pacificus*, are complicated by the fact that most research groups are usually small and have limited access to bioinformatic support to guide the analysis of whole-genome sequencing (WGS) data. Although relatively user-friendly analysis pipelines for mutant WGS data exist (Minevich *et al*. 2012; Etherington *et al*. 2014; Sun and Schneeberger 2015; Wachsman *et al*. 2017), they have been developed in species-specific manners and the integration of new organisms is often as complex as setting up a new pipeline. Thus, the research community still needs simple and well documented workflows that are composed of software packages that can easily be installed so that biologists with minimal computational skills can carry out their own analysis of WGS data. Genome sequencing of *C. elegans* and *P. pacificus* mutants often revealed in the order of thousands of candidate mutations even after successive rounds backcrossing (Rae *et al*. 2012; Iatsenko *et al*. 2013). Subsequent filtering for mutations with effects on protein sequences and additional mapping data can reduce the number of candidates by a factor of hundred, which is often sufficient for further manual inspection and experimental validation (Witte *et al*. 2015; Sieriebriennikov *et al*. 2020). Mapping data can be generated from recombinant inbred lines (Witte *et al*. 2015; Kieninger *et al*. 2016) or pooled sequencing of progeny with a mutant phenotype (Schneeberger *et al*. 2009; Doitsidou *et al*. 2010). In a previous study, we crossed a mutant line with a wildtype *P. pacificus* strain PS1843 (Washington) and sequenced a pool of progeny showing the mutant phenotype (Figure 1). The candidate region showed a pronounced bias to the reference allele. In combination with WGS of the pure mutant line, this resulted in the identification of the *P. pacificus* ortholog of *C. elegans nhr-1* as a regulator of mouthform plasticity (Sieriebriennikov *et al*. 2020).

**Figure 1.**
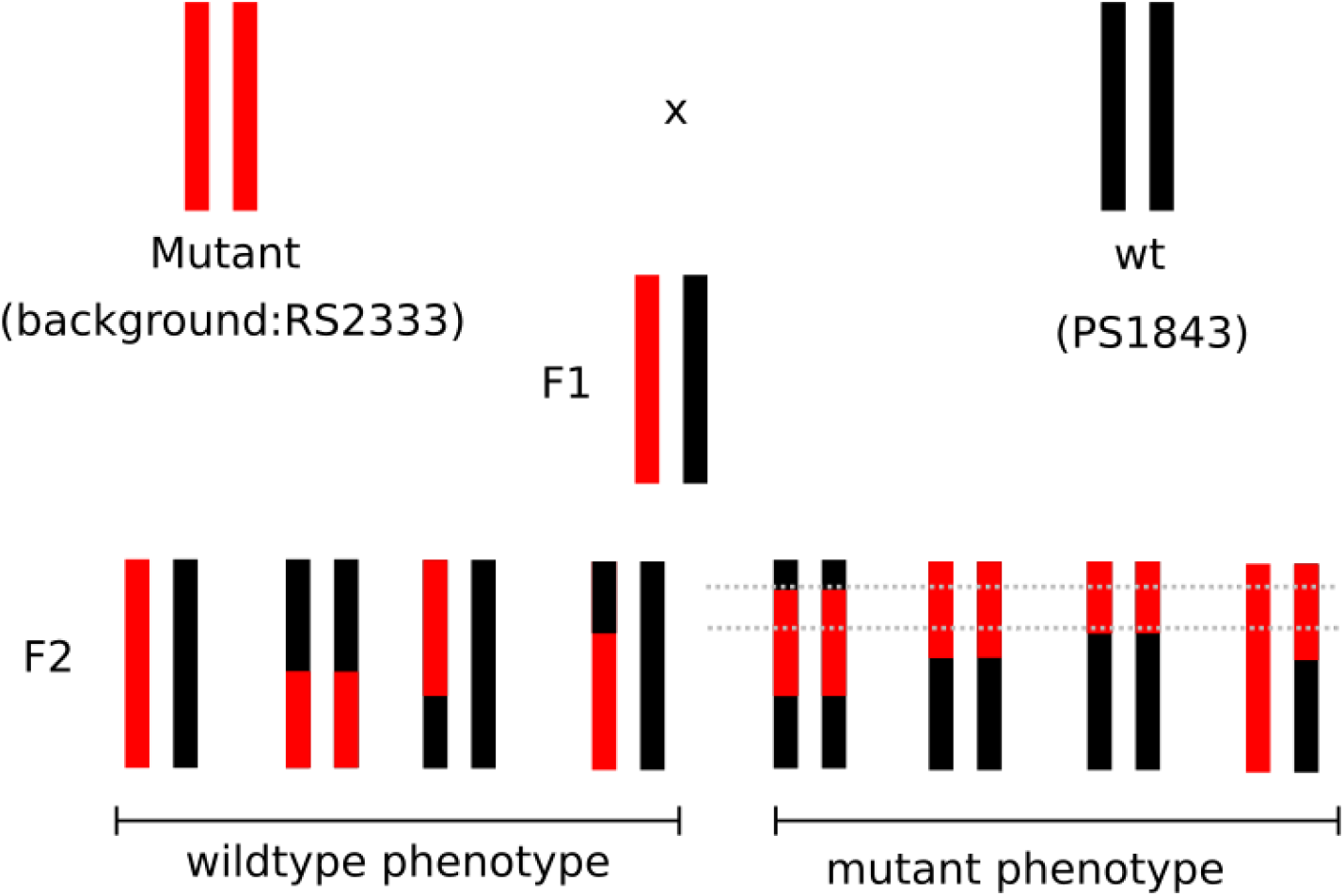
Generation of a mapping panel for pooled sequencing. A recessive mutant allele can be mapped by crossing a mutant line to a wildtype isolate and screening the F2 progeny for individuals showing the mutant phenotype. Selecting these individuals for pooled sequencing establishes a mapping panel that shows an allelic bias towards the reference alleles in the candidate region, which is highlighted by the gray dotted lines.

Here, I present a step-by-step interactive workflow for the analysis of WGS data from *P. pacificus* mutant lines with simultaneous mapping data. The effectiveness of this workflow is demonstrated by the reanalysis of the WGS data of the *P. pacificus nhr-1* mutant (Sieriebriennikov *et al*. 2020). This computational pipeline is intended as a basic workflow to generate first results, but individual components can be added or be replaced to optimize and tailor the workflow according to specific needs.

## Results and Discussion

To allow small research groups without extensive bioinformatic expertise to analyze their own genome data, I present a simplified workflow for the analysis of WGS data from *P. pacificus* mutants. The bioinformatic workflow (Figure 2) is mostly composed of widely used and well-maintained software packages such as BWA and samtools and is extended by two custom source codes for estimating the allele frequencies in pooled sequencing data (Supplemental File: Source code 1) and visualization (Supplemental File: Source code 2). The first step of the analysis consists in the alignment of the raw sequencing data to the *P. pacificus* reference assembly. This step can be executed in parallel for the WGS data from pure mutant lines, the mapping panel, and the wildtype strain PS1843 (Washington) for which WGS data was generated previously as part of population genomic studies of *P. pacificus* (Rödelsperger *et al.* 2014; McGaughran *et al*. 2016). In the second step, the alignments of the wildtype *P. pacificus* strain PS1843 serve for the identification of roughly 1,300,000 marker positions for which allele frequencies can be computed from the data of the mapping panel (Figure 2). To this end, the samtools mpileup command generates a text file which contains aligned reference and non-reference nucleotides for any given position, which is subsequently parsed by custom perl script (Supplemental File: Source code 1). This yields a data table that can be plotted with the help of R (Supplemental File: Source code 2). In the case of the mapping data of the *nhr-1* mutant line, this revealed a high frequency of reference alleles on large parts of *P. pacificus* chromosome I (5-16Mb). The causative gene, *P. pacificus nhr-1* is located at position 5.7Mb on chromosome X (Sieriebriennikov *et al*. 2020). With this candidate interval, variants of the pure mutant line can be filtered with the help of the bcftools programs. More precisely, in a first step, variants in the candidate interval are selected, subsequently background variants that are common between the *nhr-1* mutant line and its genetic background are removed (Kieninger *et al*. 2016), and the remaining variants are screened for missense mutations in protein-coding exons. In the case of the *P. pacificus nhr-1* mutant data this reduced the number of candidate mutations from roughly 3,500 single nucleotide variants to candidate mutations (Table 2). This demonstrates the effectiveness of this workflow to identify reasonably small numbers of candidate mutations after analysis of mapping data and WGS of a mutant line.

**Figure 2.**
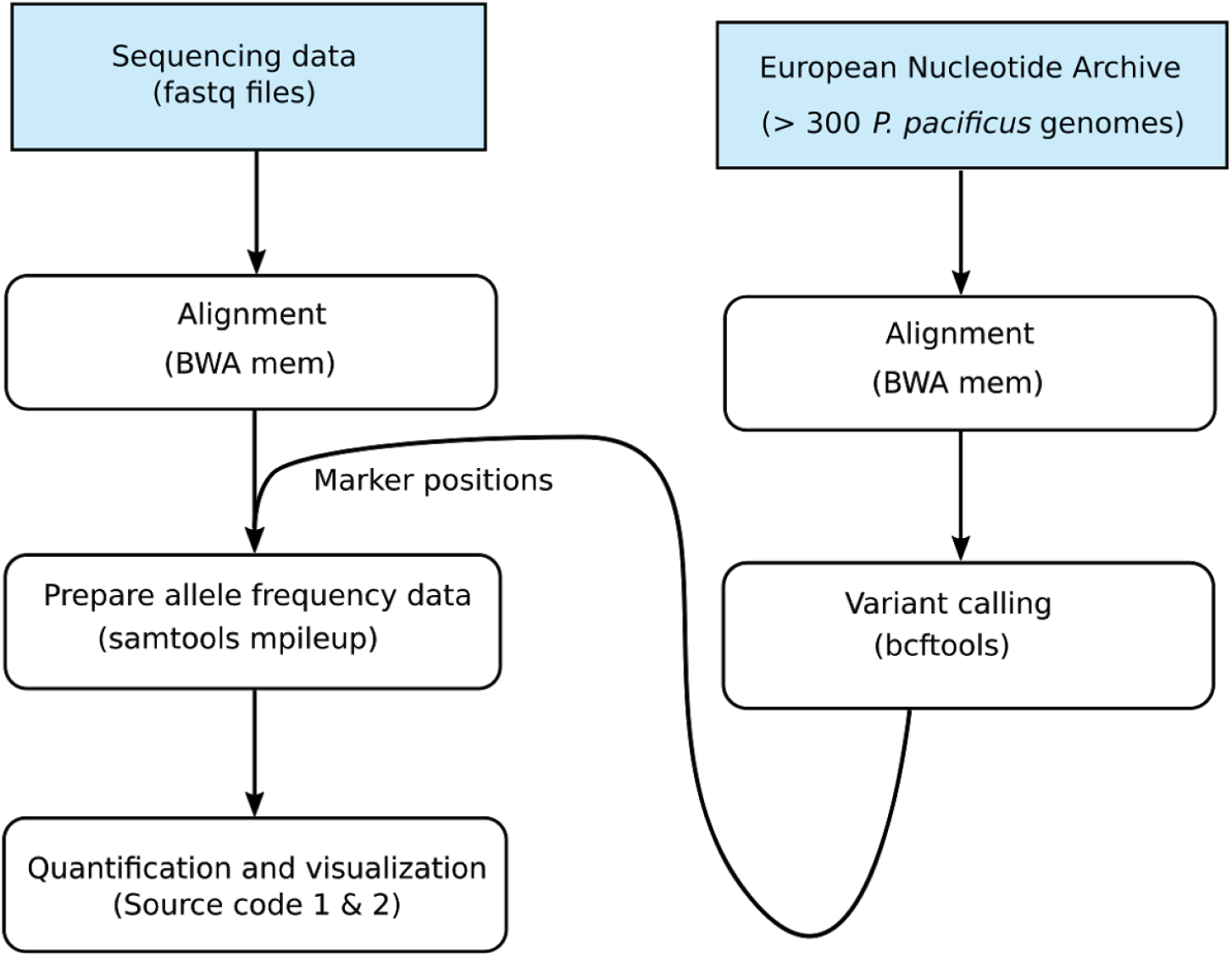
Computational workflow for mapping the candidate region. WGS data of more than 300 natural isolates of *P. pacificus* (Rödelsperger *et al*. 2014; McGaughran *et al*. 2016) can be obtained from the NCBI Sequence Read and European Nucleotide Archive. These data can be reprocessed to establish informative marker positions for the sequencing data of the pooled mapping panel. Raw read alignment is carried out by BWA and variant analysis is performed by bcftools and samtools, which are widely used and easy to install software packages for the analysis of next generation sequencing data (Li *et al*. 2009; Li and Durbin 2010; Li 2011).

The presented workflow represents a baseline approach to analyze WGS data from mutant lines. It can further be modified by adding steps for quality filtering of the raw sequencing data (Bolger *et al*. 2014), alternative variant calling procedures (McKenna *et al*. 2010), or reporting of variants in putative cis-regulatory regions that are highly conserved with other *Pristionchus* genomes (Prabh *et al*. 2018). Further, once a basic analysis pipeline is established, it can be adapted to alternative experimental designs, such as additional pooled sequencing of a control population that does not show the mutant phenotype. This may help to distinguish an association with the causative gene from segregation distortion possibly due to selfish elements or hybrid incompatibilities. In the example of the cross between the mutant strain and a wildtype strain PS1843 from Washington, there is substantial deviation from the expected 50% segregation on ChrV (Figure 3). Such segregation distortion could result in false signals if only the positive population showing the mutant phenotype is sequenced. In theory, the sequencing data of the mapping panel showing the mutant phenotype should be sufficient to identify the candidate mutations (Schneeberger *et al*. 2009; Doitsidou *et al*. 2010). However, I would still recommend to sequence the pure mutant line, because certain structural variations in the wildtype strain could inflate the number of variants in the candidate region. In addition, the sequencing data of the pure mutant line will be helpful to filter out candidates in a follow up suppressor screen. Further, if the marker density is high, restriction site associated DNA (RAD) sequencing can be applied to reduce the cost of sequencing and protocols for WGS of individual worms are available, which could provide a much better definition of the boundaries of the candidate interval. We have previously successfully applied such methods for mapping associations between genotypes and phenotypes and population genomic analysis (Witte *et al*. 2015; Falcke *et al*. 2018; Zhou *et al*. 2019). Thus, there is considerable flexibility in the design of the underlying experiments which will eventually require the computational workflow to be readjusted. Nevertheless, the presented workflow should be suitable as a baseline approach for analyzing WGS data from mutant lines.

**Figure 3.**
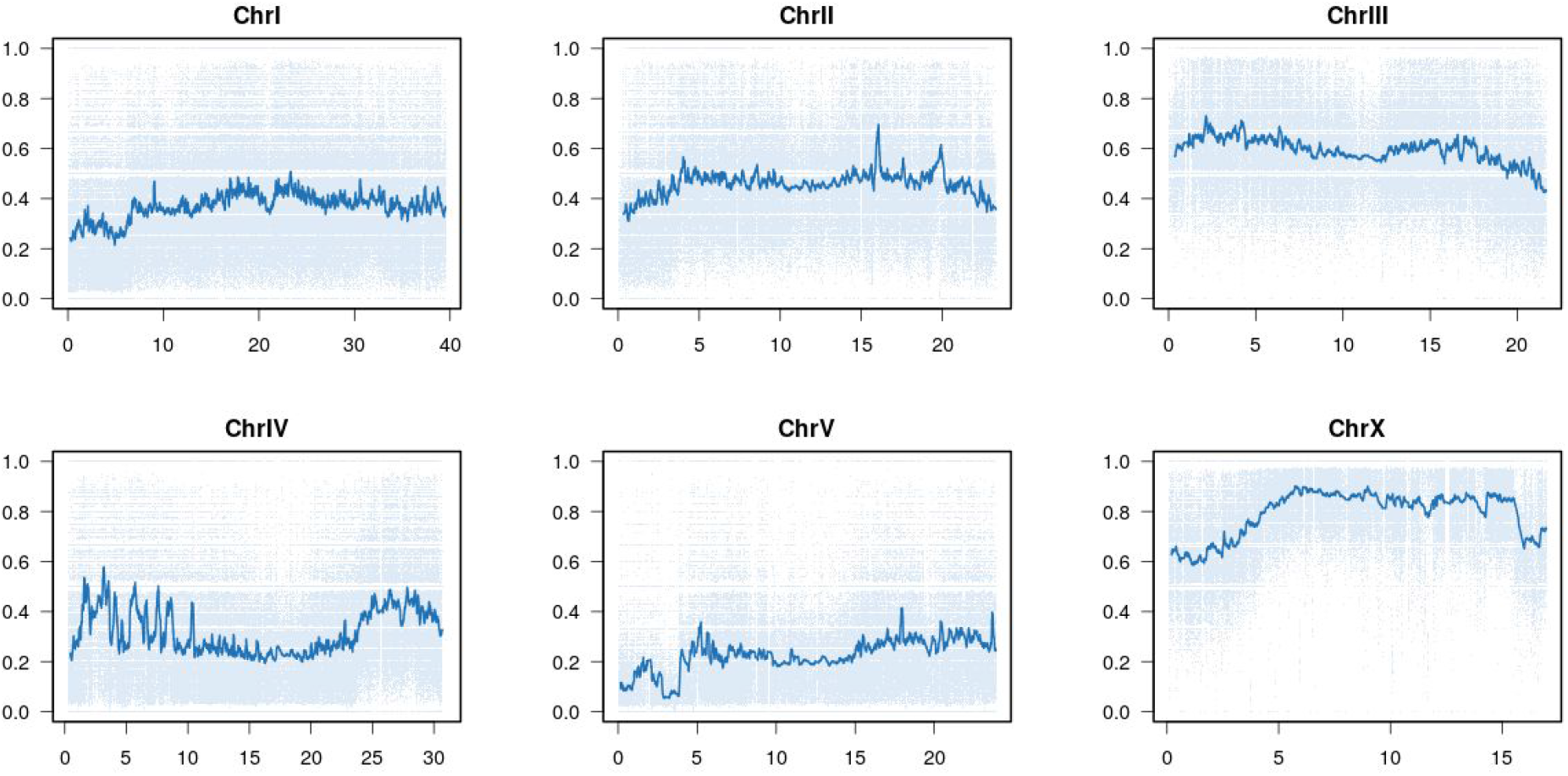
Mapping of *nhr-40* suppressor mutant. Along each *P. pacificus* chromosome, we quantified the frequency of the reference allele in the mapping panel. The x-axis denotes the Megabase positions for each chromosome and the y-axis denotes the reference allele frequency. Each marker is plotted as a light blue dot and the blue line represents a running average. The X chromosome shows the most pronounced shifts in allele-frequencies in the region 5-16Mb. The causative gene, *P. pacificus nhr-1* is located at position 5.7Mb on chromosome X.

## Methods

### Sequence retrieval, indexing of the reference genome, and alignment

The *P. pacificus* reference assembly (version El Paco (Rödelsperger *et al*. 2017)) was downloaded from http://pristionchus.org/download/. In the first step, the fasta file was decompressed and an index for efficient searching was built using the BWA program (version 0.7.17-r1188) (Li and Durbin 2009).

~~~
>bwa index genome.fa
~~~

Next, fastq files (Table 1) were downloaded from the European Nucleotide Archive, decompressed, renamed and were aligned against the reference genome with the BWA mem program.

~~~
>bwa mem -t 2 -o alignment.sam genome.fa read_1.fq read_2.fq
~~~

**Table 1.**
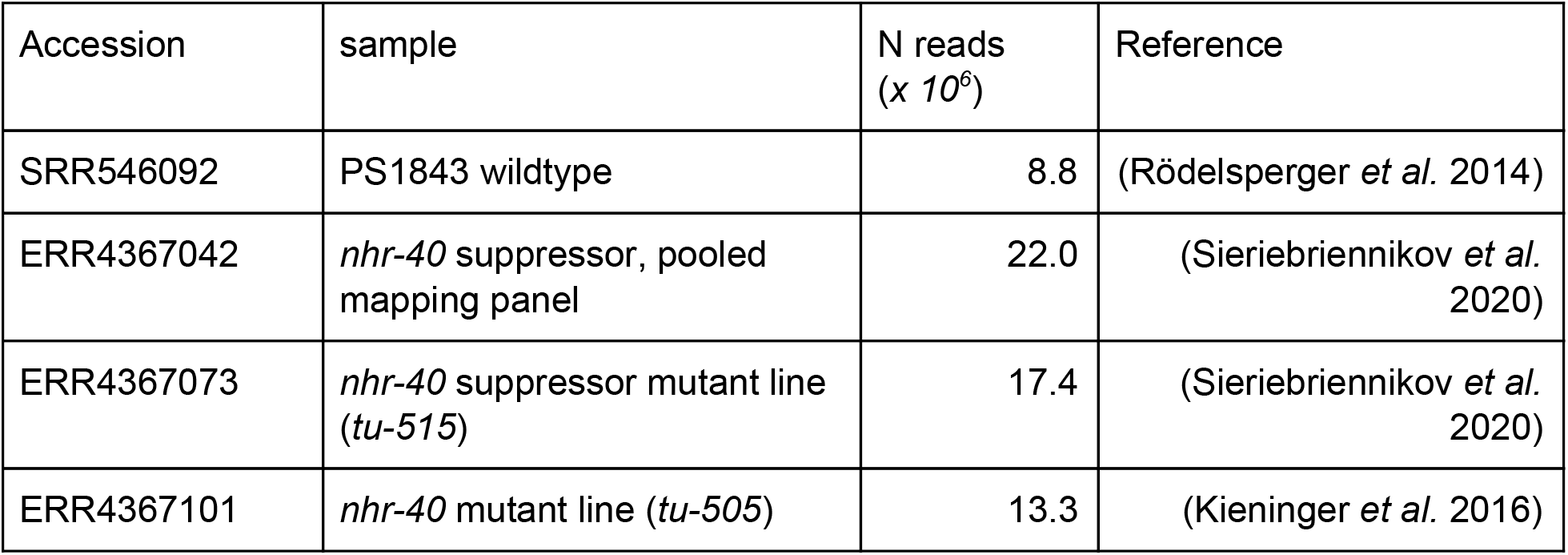
WGS data of *P. pacificus*. The table shows the run accessions from the NCBI sequence read archive and the European Nucleotide Archive for the data sets that are used in this study.

| Accession | sample | N reads<br>($\times 10^6$) | Reference |
| --- | --- | --- | --- |
| SRR546092 | PS1843 wildtype | 8.8 | (Rödelsperger *et al.* 2014) |
| ERR4367042 | *nhr-40* suppressor, pooled mapping panel | 22.0 | (Sieriebriennikov *et al.* 2020) |
| ERR4367073 | *nhr-40* suppressor mutant line (*tu-515*) | 17.4 | (Sieriebriennikov *et al.* 2020) |
| ERR4367101 | *nhr-40* mutant line (*tu-505*) | 13.3 | (Kieninger *et al.* 2016) |

The option -o specifies the name of the output file and the option -t indicates the numbers of processor cores that should be used for parallel computing. Subsequently the alignments were converted from text format (.sam) into binary format (.bam), sorted, and indexed.

~~~
>samtools view -S -b alignment.sam > alignment_unsorted.bam
>samtools sort -o alignment.bam alignment_unsorted.bam
>samtools index alignment.bam
~~~

Finally, temporary alignment files were removed to save disk space.

~~~
>rm alignment_unsorted.bam alignment.sam
~~~

### Identification of the candidate region

To identify marker positions, initial variant calls for the *P. pacificus* strains PS1843 (Washington) were generated with the program bcftools (version: 1.7) (Li 2011).

~~~
>bcftools mpileup -Ou -f genome.fa alignment.bam | bcftools call -mv -Ov -o
variants.vcf
~~~

The bcftools mpileup command calculates the genotype likelihoods that are subsequently used by the bcftools call command to generate the variant calls. The -O option specifies the output format and the -mv option indicates that only variant sites are reported. Next, the unix commands awk and grep are used to define variant positions with a quality score >100 as marker positions (indels are excluded).

~~~
>awk ‘{if($6>100) print}’ variants.vcf | grep -v INDEL |awk ‘{print $1 “\t” $2}’ >
positions.txt
~~~

These positions are then taken to generate the pileup data of all nucleotides that are aligned to a particular position.

~~~
>samtools mpileup -f genome.fa -l positions.txt mapping.bam > pileup_data.txt
~~~

Source code 1 is then copied into a text file (count_allele_frequencies.pl) and is used to count the allele frequencies for each position in the pileup data.

~~~
>perl count_allele_frequencies.pl pileup_data.txt > mapping_data.txt
~~~

Within the same directory, source code 2 has to be executed within R to generate the plot of allele frequency data.

### Identification of candidate mutations

Initial variant calling for the alignments of the *nhr-1* mutant line and other samples are generated by bcftools.

~~~
>bcftools mpileup -Ou -f genome.fa alignment.bam | bcftools call -mv -Ou | bcftools
view -i ‘%QUAL>=20’ -Oz > variants.vcf.gz
>bcftools index variants.vcf.gz
~~~

Here, the additional call of the bcftools view command filters for variants with a quality score > 20. The bcftools index command is needed for the following filtering procedures. Next, variants falling into the candidate region are extracted and the resulting compressed .vcf file is indexed to allow further filtering.

~~~
>bcftools filter -O z -r ChrX:5000000-16000000 variants.vcf.gz > filtered1.vcf.gz
>bcftools index filtered1.vcf.gz
~~~

The following steps extracts all variant positions that are shared with the genetic background (*nhr-40* mutant line (Kieninger *et al*. 2016)) and the corresponding variants are extracted.

~~~
>bcftools isec -C filtered1.vcf.gz nhr-40.vcf.gz > pos.txt
>bcftools filter -O z -R pos.txt filtered1.vcf.gz > filtered2.vcf.gz
>bcftools sort -O z filtered2.vcf.gz > filtered2_sorted.vcf.gz
>bcftools index filtered2_sorted.vcf.gz
~~~

Finally, the program bcftools csq can be used for functional annotation of the filtered variants. To this end, the bcftools compatible version of the *P. pacificus* gene annotation (version: El Paco annotations v2.1 (Rödelsperger *et al*. 2019)) was downloaded from http://www.pristionchus.org/download/.

~~~
>bcftools csq -p s -f genome.fa -g El_Paco_annotation_v2_1_bcftools_compatible.gff3
filtered2_sorted.vcf.gz > variants_annotated.vcf
~~~

Filtering the annotated variant file for missense mutations identifies ten candidate mutations (Table 2) including the non-synonymous mutation in the *P. pacificus nhr-1* gene (Sieriebriennikov *et al*. 2020).

~~~
>grep missense variants_annotated.vcf
~~~

**Table 2.** Final candidate list of mutations. This list contains all mutations in the candidate interval that are classified to change protein-coding genes and are not found in the sequencing data of the genetic background (Kieninger *et al*. 2016).

| Position | Substitution | *P. pacificus* gene ID /<br>Amino acid substitution | *C. elegans*<br>ortholog |
| --- | --- | --- | --- |
| ChrX:5753767 | C>T | ppa_stranded_DN28508_c0_g1_i2 /<br>158R>158W | *nhr-1* |
| ChrX:6957752 | C>T | ppa_stranded_DN25215_c0_g1_i1 /<br>186D>186N | *sec-3* |
| ChrX:7145062 | C>T | PPA39440 / 296D>296N | B0361.3 |
| ChrX:8866724 | C>T | PPA39709 / 240E>240K | - |
| ChrX:9387377 | C>T | PPA07538 / 419D>419N | *flp-11* |
| ChrX:9894810 | G>A | PPA20385 / 393T>393I | - |
| ChrX:13784115 | A>T | PPA25760 / 438S>438 | - |
| ChrX:13855207 | G>A | PPA40003 / 211D>211N | - |
| ChrX:13882049 | G>A | PPA40012 / 96T>96I | *srh-50* |
| ChrX:14602500 | C>T | PPA42301 / 396R>396C | - |

## Acknowledgements

I would like to thank the whole *Pristionchus* community for their long-term interest in studying *P. pacificus* and thus motivating this work. Special thanks goes to Erick Rios and Iris Rafaela for testing this workflow and helpful comments on this manuscript.

## Supplementary Material

**File name:** Supplemental_files.zip

The Supplemental file is a zip archive containing extended documentation, source codes and example files.

