## Supplementary material for "A simplified workflow for the analysis of whole-genome sequencing data from *Pristionchus pacificus* mutant lines": Extended_documentation_source_code2.txt

Content:

Extended_documentation_source_code2.txt   -  this file
mapping_data.txt   - downsampled mapping data for testing source code 2
variants.vcf.gz    - variant file for the nhr-40 suppressor mutant line
nhr-40.vcf.gz      - variant file for the nhr-40 background
source_code1.pl    - custom perlscript


#################################################################
#                                                               #
# 1   Raw read alignment with BWA                               #
#                                                               #
#################################################################


1.1 Test if BWA is installed

To test if the software suite BWA is installed on your computer, enter
the command line and type: 

>bwa 

If the program is installed, the program will output a general help message
that specifies the version and an overview of different BWA programs.


Program: bwa (alignment via Burrows-Wheeler transformation)
Version: 0.7.17-r1188
Contact: Heng Li <>

Usage:   bwa <command> [options]

Command: index         index sequences in the FASTA format
         mem           BWA-MEM algorithm
         fastmap       identify super-maximal exact matches
         pemerge       merge overlapping paired ends (EXPERIMENTAL)
         aln           gapped/ungapped alignment
         samse         generate alignment (single ended)
         sampe         generate alignment (paired ended)
         bwasw         BWA-SW for long queries

         shm           manage indices in shared memory
         fa2pac        convert FASTA to PAC format
         pac2bwt       generate BWT from PAC
         pac2bwtgen    alternative algorithm for generating BWT
         bwtupdate     update .bwt to the new format
         bwt2sa        generate SA from BWT and Occ

Note: To use BWA, you need to first index the genome with `bwa index'.
      There are three alignment algorithms in BWA: `mem', `bwasw', and
      `aln/samse/sampe'. If you are not sure which to use, try `bwa mem'
      first. Please `man ./bwa.1' for the manual.

If BWA is not installed, visit https://github.com/lh3/bwa
and follow the documentation for installation. On Ubuntu Linux operating systems,
the packages BWA, samtools, and bcftools can be installed using the standard
package management systems such as apt or aptitude.

1.2 Indexing the genome

The most recent assembly of the Prisitonchus pacificus genome can
be retrieved here: 

http://pristionchus.org/download/El_Paco_genome.tgz

This compressed archive can be downloaded and expanded by the command:

>tar -xzvf El_Paco_genome.tgz

The archive contains a fasta file El_Paco_genome.fa, which can be indexed
using the bwa index command

bwa index El_Paco_genome.fa

This will run for a couple of minutes and create a number of files which
represent the indexed genome and allow the BWA program to search efficiently
for sequence matches.

El_Paco_genome.fa.amb
El_Paco_genome.fa.ann
El_Paco_genome.fa.bwt
El_Paco_genome.fa.pac
El_Paco_genome.fa.sa

The output of the bwa index command will look like this:

[bwa_index] Pack FASTA... 1.22 sec
[bwa_index] Construct BWT for the packed sequence...
[BWTIncCreate] textLength=317000792, availableWord=34305004
[BWTIncConstructFromPacked] 10 iterations done. 56587912 characters processed.
[BWTIncConstructFromPacked] 20 iterations done. 104542152 characters processed.
[BWTIncConstructFromPacked] 30 iterations done. 147159992 characters processed.
[BWTIncConstructFromPacked] 40 iterations done. 185034776 characters processed.
[BWTIncConstructFromPacked] 50 iterations done. 218693944 characters processed.
[BWTIncConstructFromPacked] 60 iterations done. 248606280 characters processed.
[BWTIncConstructFromPacked] 70 iterations done. 275188424 characters processed.
[BWTIncConstructFromPacked] 80 iterations done. 298810664 characters processed.
[bwt_gen] Finished constructing BWT in 89 iterations.
[bwa_index] 84.24 seconds elapse.
[bwa_index] Update BWT... 0.72 sec
[bwa_index] Pack forward-only FASTA... 0.60 sec
[bwa_index] Construct SA from BWT and Occ... 39.08 sec
[main] Version: 0.7.17-r1188
[main] CMD: bwa index El_Paco_genome.fa
[main] Real time: 132.965 sec; CPU: 125.861 sec


In the following, it is assumed that all analysis is carried out in the
same folder that contains the indexed genome


1.3 Obtaining Fastq files with sequencing data  

Visit the European Nucleotide Archive

https://www.ebi.ac.uk/ena/

and type into the search field one of the accessions from Table 1, e.g. SRR546092 or ERR4367073.
Click on the link under the Run menu and download the compressed fastq files (.fastq.gz) corresponding
to the forward and reverse reads, e.g.

ERR4367073_1.fastq.gz
ERR4367073_2.fastq.gz

Uncompress these files in the same folder that contains the indexed genome.

>gzip -d ERR4367073_1.fastq.gz
>gzip -d ERR4367073_2.fastq.gz

This should be done for all samples in table 1.


1.4 Align raw reads to the Pristionchus pacificus genome


>bwa mem -t 2 El_Paco_genome.fa ERR4367073_1.fastq ERR4367073_2.fastq > nhr-1.sam
>bwa mem -t 2 El_Paco_genome.fa ERR4367101_1.fastq ERR4367101_2.fastq > nhr-40.sam
>bwa mem -t 2 El_Paco_genome.fa ERR4367042_1.fastq ERR4367042_2.fastq > mapping.sam
>bwa mem -t 2 El_Paco_genome.fa SRR546092_1.fastq  SRR546092_2.fastq > PS1843.sam

Aligning a single data set can take up to hours, during the computation the program
output looks like this.

[M::bwa_idx_load_from_disk] read 0 ALT contigs
[M::process] read 99010 sequences (10000010 bp)...
[M::process] read 99010 sequences (10000010 bp)...
[M::mem_pestat] # candidate unique pairs for (FF, FR, RF, RR): (8, 38296, 1, 1)
[M::mem_pestat] skip orientation FF as there are not enough pairs
[M::mem_pestat] analyzing insert size distribution for orientation FR...
[M::mem_pestat] (25, 50, 75) percentile: (195, 210, 224)
[M::mem_pestat] low and high boundaries for computing mean and std.dev: (137, 282)
[M::mem_pestat] mean and std.dev: (209.32, 22.44)
[M::mem_pestat] low and high boundaries for proper pairs: (108, 311)
[M::mem_pestat] skip orientation RF as there are not enough pairs
[M::mem_pestat] skip orientation RR as there are not enough pairs
[M::mem_process_seqs] Processed 99010 reads in 12.695 CPU sec, 12.627 real sec
[M::process] read 99010 sequences (10000010 bp)...
[M::mem_pestat] # candidate unique pairs for (FF, FR, RF, RR): (4, 38277, 0, 0)
[M::mem_pestat] skip orientation FF as there are not enough pairs
[M::mem_pestat] analyzing insert size distribution for orientation FR...
[M::mem_pestat] (25, 50, 75) percentile: (195, 209, 224)
[M::mem_pestat] low and high boundaries for computing mean and std.dev: (137, 282)
[M::mem_pestat] mean and std.dev: (209.38, 22.19)
[M::mem_pestat] low and high boundaries for proper pairs: (108, 311)
[M::mem_pestat] skip orientation RF as there are not enough pairs
[M::mem_pestat] skip orientation RR as there are not enough pairs
[M::mem_process_seqs] Processed 99010 reads in 14.021 CPU sec, 13.942 real sec
[M::process] read 99010 sequences (10000010 bp)...

....

[M::process] read 86694 sequences (8756094 bp)...
[M::mem_pestat] # candidate unique pairs for (FF, FR, RF, RR): (7, 38472, 0, 1)
[M::mem_pestat] skip orientation FF as there are not enough pairs
[M::mem_pestat] analyzing insert size distribution for orientation FR...
[M::mem_pestat] (25, 50, 75) percentile: (195, 209, 224)
[M::mem_pestat] low and high boundaries for computing mean and std.dev: (137, 282)
[M::mem_pestat] mean and std.dev: (209.15, 22.21)
[M::mem_pestat] low and high boundaries for proper pairs: (108, 311)
[M::mem_pestat] skip orientation RF as there are not enough pairs
[M::mem_pestat] skip orientation RR as there are not enough pairs
[M::mem_process_seqs] Processed 99010 reads in 13.561 CPU sec, 13.469 real sec
[M::mem_pestat] # candidate unique pairs for (FF, FR, RF, RR): (4, 33519, 0, 2)
[M::mem_pestat] skip orientation FF as there are not enough pairs
[M::mem_pestat] analyzing insert size distribution for orientation FR...
[M::mem_pestat] (25, 50, 75) percentile: (196, 209, 224)
[M::mem_pestat] low and high boundaries for computing mean and std.dev: (140, 280)
[M::mem_pestat] mean and std.dev: (209.65, 21.75)
[M::mem_pestat] low and high boundaries for proper pairs: (112, 308)
[M::mem_pestat] skip orientation RF as there are not enough pairs
[M::mem_pestat] skip orientation RR as there are not enough pairs
[M::mem_process_seqs] Processed 86694 reads in 12.558 CPU sec, 12.519 real sec
[main] Version: 0.7.17-r1188
[main] CMD: bwa mem -t 1 ppac.contigs-1.04.fasta RS2333_1.fq RS2333_2.fq
[main] Real time: 4078.348 sec; CPU: 4105.139 sec


#################################################################
#                                                               #
# 2   Converting the alignments into .bam format                #
#                                                               #
#################################################################

The following commands convert the .sam output files that were
generated by BWA into binary alignment (.bam) files. These files
consume less disk space and can be more efficiently handled by the
downstream programs. Each of the following commands can take up to
several minutes on a standard Desktop computer.

2.1 Conversion from text (.sam) into binary (.bam) format 

>samtools view -S -b PS1843.sam  > PS1843_unsorted.bam
>samtools view -S -b mapping.sam > mapping_unsorted.bam
>samtools view -S -b nhr-1.sam  > nhr-40_unsorted.bam
>samtools view -S -b nhr-40.sam > nhr-40_unsorted.bam


2.2 Sorting the binary slignment files

>samtools sort -o mapping.bam -@ 2 mapping_unsorted.bam
>samtools sort -o PS1843.bam -@ 2  PS1843_unsorted.bam 
>samtools sort -o nhr-1.bam -@ 2  nhr-1_unsorted.bam 
>samtools sort -o nhr-40.bam -@ 2  nhr-40_unsorted.bam 

2.3 Indexing the binary slignment files

>samtools index PS1843.bam
>samtools index nhr-1.bam
>samtools index nhr-40.bam
>samtools index mapping.bam

The last command will generate index files with the file extension .bam.bai.
If everything worked fine, the resulting .bam and .bam.bai. files should
have the following file sizes.

>ls -lh *.bam*
 
3,2G mapping.bam
466K mapping.bam.bai
2,2G nhr-1.bam
462K nhr-1.bam.bai
1,9G nhr-40.bam
460K nhr-40.bam.bai
1,5G PS1843.bam
442K PS1843.bam.bai

The temporary  _unsorted.bam and .sam files can now be deleted:

>rm *_unsorted.bam *.sam


##################################################################
#                                                                #
# 3   Initial variant calling and mapping the candidate region   #
#                                                                #
##################################################################

3.1 Initial variant calling can take several minutes for each indiviudal data set.

>bcftools mpileup -Ou -f El_Paco_genome.fa  PS1843.bam | bcftools call -mv -Ov -o PS1843.vcf

The following steps call variants of the previously generated nhr-40 data set and prepare
it for filtering (section 4)

>bcftools mpileup -Ou -f El_Paco_genome.fa  nhr-40.bam | bcftools call -mv -Ov -o nhr-40.vcf
>bgzip nhr-40.vcf
>bcftools index nhr-40.vcf.gz

3.2 Calculation allele frequencies at marker positions 

To identify suitable markers for the bulked-segregant analysis, I filter high quality
variant positions from the resulting vcf file:

>awk '{if($6>100) print}' PS1843.vcf  | grep -v INDEL |awk '{print $1 "\t" $2}'  > positions.txt 

Next, the mpileup files will be generated for the resulting 1.3 million variant positions. The
mpileup data contains the inforamation of every nucleotide from different reads that are aligned
at a given position.

>samtools mpileup  -f El_Paco_genome.fa  -l positions.txt  mapping.bam  > mapping_raw_pileup.txt

In the next step, the perl script allele_counts.pl (Source code 1) is run to extract the allele
frequency data from the mpileup files.

>perl source_code1.pl mapping_raw_pileup.txt  > mapping_data.txt 


3.3 Generate the mapping plot

Enter the R environment

>R

Copy and paste the following code into the R shell


## Source code 2
## Copy from here

running_mean <- function(y, n=100){
  y.new = c()
  for(i in n:length(y) ){
    y.new[i] = mean(y[(i-n+1):(i+1)])
  }
  y.new
}

A <- read.table("mapping_data.txt")  ## input data
A <- subset( A, A[,3] + A[,4] > 20 )

chromos <-  c("ChrI", "ChrII", "ChrIII", "ChrIV", "ChrV", "ChrX")

png("california_ratio.png", width=1200, height=600 )
par(mfrow=c(2,3),cex=1.2, mar=c(4,4,2,2) , lwd=2 , las=1 )  ## specify to plot 2 rows with 3 panels each

for(i in 1:6 ){   ## foreach chromosome
 
  tmp <- subset(A, A[,1] == chromos[i] ) ## select data for specific chromosome
  tmp <- tmp[order(tmp[,2]),]  ## sort by position
  x <- tmp[,2] / 1000000
  y <- tmp[,3]/(tmp[,3]+ tmp[,4])  ## reference ratio
  plot(x, y , pch=".", col=blues9[2], ylim=c(0,1) , ylab="", xlab="",   main=chromos[i])
  lines(x, running_mean(y, n=1000),  col=blues9[7] )
}
dev.off()

### Until here

This will generate a png file with the plot

Exit the R shell by typing

>q()


##################################################################
#                                                                #
# 4   Filtering of candidate variants                            #
#                                                                #
##################################################################


Initial variant calling for the alignments of the nhr-1 mutant line and other samples are generated by bcftools.
This can take several minutes.

>bcftools mpileup -Ou -f El_Paco_genome.fa nhr-1.bam | bcftools call -mv -Ou |  bcftools view -i '%QUAL>=20' -Oz >  variants.vcf.gz
>bcftools index  variants.vcf.gz

Here, the additional call of the bcftools view command filters for variants with a quality score > 20.
The bcftools index command is needed for the following filtering procedures. Next, variants falling
into the candidate region are extracted and the resulting compressed .vcf file is indexed to allow
further filtering.

>bcftools filter -O z -r ChrX:5000000-16000000 variants.vcf.gz   > filtered1.vcf.gz
>bcftools index  filtered1.vcf.gz

The following steps extracts all variant positions that are shared with the genetic background
(nhr-40 mutant line (Kieninger et al. 2016)) and the corresponding variants are extracted.


>bcftools isec -C filtered1.vcf.gz  nhr-40.vcf.gz > pos.txt
>bcftools filter -O z -R pos.txt  filtered1.vcf.gz  > filtered2.vcf.gz
>bcftools sort  -O z filtered2.vcf.gz > filtered2_sorted.vcf.gz
>bcftools index  filtered2_sorted.vcf.gz

Finally, the program bcftools csq can be used for functional annotation of the filtered variants.
To this end, the bcftools compatible version of the P. pacificus gene annotation
(version: El Paco annotations v2.1 (Rödelsperger et al. 2019)) was downloaded
from http://www.pristionchus.org/download/. 

>bcftools csq -p s -f  El_Paco_genome.fa -g  El_Paco_annotation_v2_1_bcftools_compatible.gff3 filtered2_sorted.vcf.gz > variants_annotated.vcf

Filtering the annotated variant file for missense mutations identifies ten candidate mutations (Table 2)
including the non-synonymous mutation in the P. pacificus nhr-1 gene (Sieriebriennikov et al. 2020).

>grep missense variants_annotated.vcf
