## Supplementary material for "A simplified workflow for the analysis of whole-genome sequencing data from *Pristionchus pacificus* mutant lines": source_code1.pl

```
#!/usr/bin/perl

use strict;

die "program mpileup.txt\n" if( @ARGV < 1);

open( IN, $ARGV[0] );
while(<IN>){
    my ($chr,$pos,$ref,$depth,$string) = split();
    next if( $string =~ /-/  );  ## ignore deletions
    next if( $string =~ /\+/ );  ## ignore insertions
    $string =~ s/\$//g; ## remove special symbol “$”:= last base of read
    $string =~ s/\^\]//g; ## remove  “^]” := first base of a read
    my $n = length($string);
    $string =~ tr/\.,//d;
    my $n_non_ref = length($string);
    my $n_ref     = $n - $n_non_ref;
    print "$chr\t$pos\t$n_ref\t$n_non_ref\n";
}
close( IN );


sub compare_pos{   ## this subroutine is needed to sort by genomic position
    my ($a, $b) = @_;
    my ($chr1, $p1 ) = split(/:/, $a);
    my ($chr2, $p2 ) = split(/:/, $b );
    if( $chr1 eq $chr2 ){
    return ($p1 <=> $p2 );
    }
    else{
    return ( $chr1 cmp $chr2 );
    }
}
```
